# Weather explains inter-annual variability, but not the temporal decline, in insect biomass

**DOI:** 10.1101/2024.05.02.591618

**Authors:** François Duchenne, Colin Fontaine

**Affiliations:** Swiss Federal Institute for Forest, Snow and Landscape Research (WSL), 8903 Birmensdorf, Switzerland; Estación Biológica de Doñana (EBD-CSIC), Sevilla, Spain; Centre d’Écologie et des Sciences de la Conservation (UMR 7204) - Muséum national d’Histoire naturelle, CNRS, Sorbone Université - 43, rue Buffon 75005 Paris, France

## Abstract

In a recent publication, Müller *et al*. (2023) re-analysed, in light of new data, the dataset of the highly cited paper of Hallmann *et al*. (2017) showing a strong decline in insect biomass in Germany between 1989 and 2016. In their re-analysis, Müller *et al*. completed Hallmann *et al*’s model with a focus on modelling the effects of weather conditions on insect biomass. They also included temporal changes in habitat as additional predictors, using the same variables as Hallmann *et al*., which although not entirely satisfactory due to the scarcity of historical habitat data, represent the best available data. While they trained their model on the Hallmann *et al*.’s dataset, Müller *et al*. validated it with an independent dataset. These upgraded analyses are a nice demonstration of the strong impact of climatic conditions on annual insect biomass. However, Müller *et al*. conclusion that “temporal variation in weather conditions explained most of the temporal changes in insect biomass whereas temporal changes in habitat conditions played only a minor role” was overstated. Here we argue that their methodological approach was unsuitable to draw such conclusion, because of omitted variable bias. We show that more appropriate analyses produce a pattern opposite to the main conclusion of Müller *et al*.: there is a significant temporal decline in insect biomass not explained by weather conditions.

## Main

In a recent publication^1^, Müller *et al*. re-analysed, in light of new data, the dataset of the highly cited paper of Hallmann *et al*.^2^ showing a strong decline in insect biomass in Germany between 1989 and 2016. In their re-analysis, Müller *et al*. completed Hallmann et al’s model with a focus on modelling the effects of weather conditions on insect biomass. They also included temporal changes in habitat as additional predictors, using the same variables as Hallmann *et al*., which although not entirely satisfactory due to the scarcity of historical habitat data, represent the best available data. While they trained their model on the Hallmann *et al*.’s dataset, Müller *et al*. validated it with an independent dataset. These upgraded analyses are a nice demonstration of the strong impact of climatic conditions on annual insect biomass.

Müller *et al*. concluded that the observed decline and rise of insect biomass, were mostly explained by variations in weather conditions. Here we argue that their analysis was unsuitable to draw such conclusion, because it left out alternative potential drivers of insect biomass, which should be incorporated into the statistical model before one can conclude on the main causes of temporal variations in insect biomass. Omitted drivers, such as pesticides, and complex landscape-level habitat change, are likely to be correlated over time with the main studied driver (weather conditions), because they are all affected by anthropogenic drivers (Fig. 1a). Neglecting possible confounders might lead to bias, which is sometimes known as the Omitted Variable Bias^3,4^. Here, we show that more appropriate analyses produce a pattern opposite to the main conclusion of Müller *et al*.: there is a significant temporal decline in insect biomass not explained by weather conditions.

**Fig. 1:**
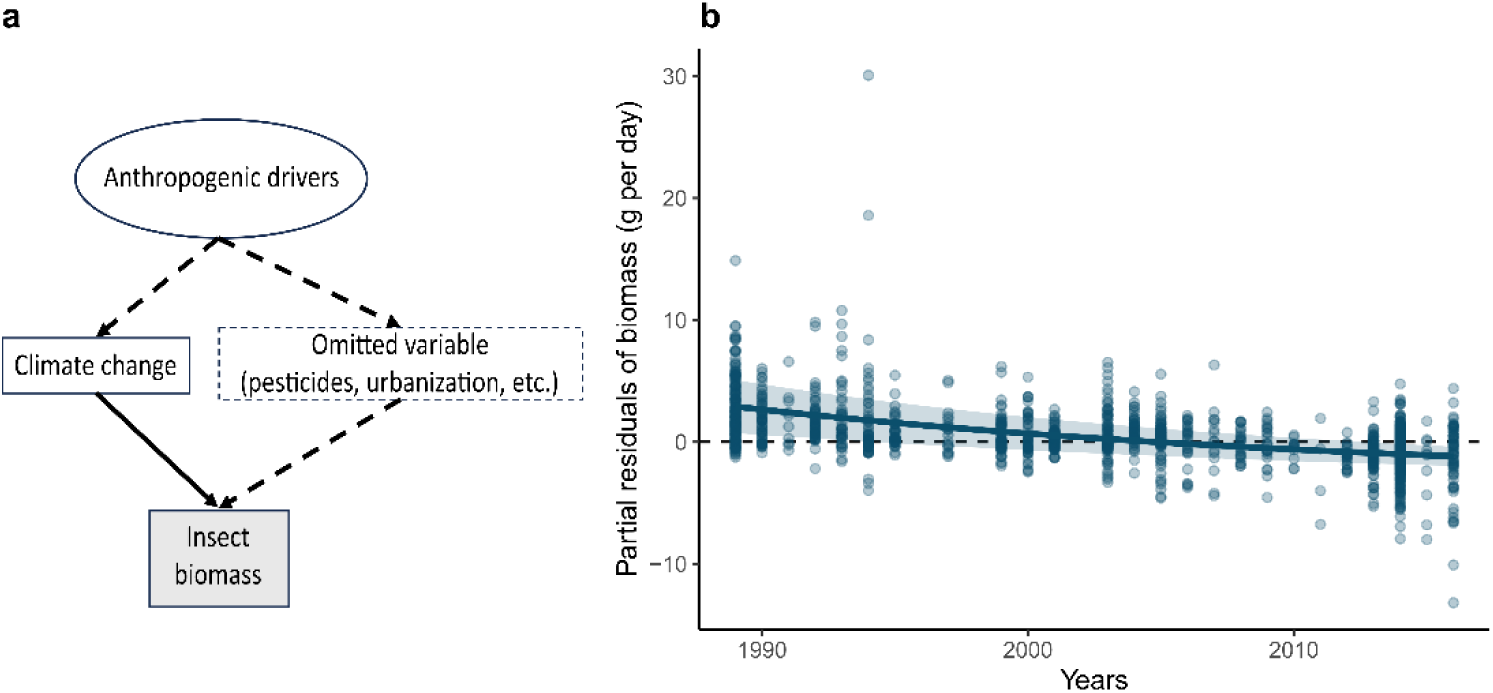
The temporal trend in insect biomass is significantly negative when the effects of weather are accounted for. (a) illustration of a possible omitted variable (dashed white box), linked to the studied variables (climate change) because it shares common drivers (ellipse), of insect biomass (response variable, grey box). Dashed arrows show the unmodelled linked, while solid arrow shows the linked studied by Müller et al. (b) The partial residuals of biomass, i.e. the amount of biomass not explained by other predictors than year, as a function of year. Line and ribbon show the model prediction and its 95% confidence interval, respectively.

### A strong temporal decline in insect biomass not explained by weather conditions

Müller *et al*. argue that weather conditions were the only important driver of temporal changes in insect biomass, because when weather conditions were included as explanatory variables, the model residuals exhibited no temporal trend (model 5 of their study). Estimating the temporal trend in residuals is a hierarchical approach prone to bias because current climate change is characterized, among others, by a known temporal trend in weather conditions. The statistical fit, which seeks to explain as much variance as possible with the variables included in the model, is likely to attribute any temporal change in insect biomass to temporal changes in weather conditions. Thus, the absence of temporal trend in the model residuals is not informative on the importance of non-modelled drivers.

For example, agriculture intensification is a likely driver of insect biomass, especially pesticide use^5–7^, but neither Hallmann *et al*. nor Müller *et al*. could model its effect because spatio-temporal metrics are missing^2^. However, we know that, the total applied pesticide toxicity in Germany exhibits a strong temporal and spatial variations^8^, which can lead to confounding effects with climate change. However, since data on these potential drivers over time and space are missing, they cannot be included. While adding a random year slope do this and can be used, will fail to solve an omitted variable bias if the missing drivers are correlated to the studied ones^4^, which is likely the case. To account for potential driver that would be linked to global change, and thus exhibiting a temporal trend, we propose to estimate a temporal trend, simultaneously to other effects, to decrease the putative bias due to any temporal confounders. This would lead to add a fixed linear year effect to the model of Müller *et al*., which will compete with other effects and absorb what correlates more with time than with other drivers included in the model.

Adding a linear year effect to the covariates included in Müller *et al*.’s model, indicates that there is a significant decline in insect biomass over time (-4.0%.year^-1^) that is not explained by weather conditions (Fig. 1b and Table 1), while improving the fit of the model (lower AIC, Table 1). This temporal trend is not informative of the possible drivers of the temporal decline but indicates that insect biomass declined by 4% per year because of unknown factors.

**Table 1:** Model estimates and goodness of fit for the model of Müller et al. (2023) and for the modified version, with an additional linear effect of year.

| Variable | Model 5 from Müller *et al.* |  |  | Model 5 + year |  |  |
| --- | --- | --- | --- | --- | --- | --- |
|  | Estimate | Stde | p-value | Estimate | Stde | p-value |
| Number of herb species | 0.0008 | 0.0011 | 0.4763 | -0.0022 | 0.0011 | **0.0377** |
| Number of tree species | 0.1174 | 0.0121 | **0.0000** | 0.0515 | 0.0143 | **0.0003** |
| Ellenberg value light | 0.1469 | 0.0646 | **0.0232** | 0.0529 | 0.0635 | 0.4051 |
| Ellenberg value temperature | -0.0351 | 0.0406 | 0.3867 | 0.0702 | 0.0408 | 0.0857 |
| Proportion of arable land | -0.3530 | 0.1108 | **0.0015** | -0.0808 | 0.1130 | 0.4746 |
| Proportion of forest | -0.1493 | 0.1139 | 0.1899 | 0.0630 | 0.1139 | 0.5803 |
| Proportion of grassland | 0.3484 | 0.1161 | **0.0027** | 0.2487 | 0.1129 | **0.0277** |
| Proportion of water | 0.2816 | 0.1479 | 0.0571 | 0.0364 | 0.1448 | 0.8016 |
| *T* | 0.0814 | 0.0062 | **0.0000** | 0.0844 | 0.0060 | **0.0000** |
| *P* | -0.0033 | 0.0008 | **0.0000** | -0.0025 | 0.0007 | **0.0007** |
| *T* × *P* | -0.0001 | 0.0002 | 0.7482 | 0.0000 | 0.0002 | 0.8560 |
| *T* ano. winter | -0.2943 | 0.0268 | **0.0000** | -0.1232 | 0.0321 | **0.0001** |
| *P* ano. winter | 0.0339 | 0.0026 | **0.0000** | 0.0197 | 0.0030 | **0.0000** |
| *T* ano. winter × *P* ano. winter | -0.0114 | 0.0025 | **0.0000** | -0.0021 | 0.0026 | 0.4187 |
| *T* ano. April cur | 0.0820 | 0.0261 | **0.0017** | 0.0810 | 0.0237 | **0.0007** |
| *P* ano. April cur | 0.0148 | 0.0016 | **0.0000** | 0.0068 | 0.0017 | **0.0000** |
| *T* ano. April cur × *P* ano. April cur | -0.0028 | 0.0009 | **0.0036** | -0.0003 | 0.0009 | 0.7761 |
| *T* ano. April prev. | -0.1082 | 0.0303 | **0.0004** | 0.0155 | 0.0301 | 0.6073 |
| *P* ano. April prev. | 0.0021 | 0.0015 | 0.1477 | 0.0028 | 0.0014 | **0.0405** |
| *T* ano. April prev. × *P* ano. April prev. | -0.0044 | 0.0008 | **0.0000** | -0.0014 | 0.0008 | 0.0932 |
| *T* ano. month prev. | -0.0078 | 0.0119 | 0.5135 | -0.0059 | 0.0117 | 0.6134 |
| *P* ano. month prev. | -0.0009 | 0.0004 | **0.0369** | -0.0001 | 0.0004 | 0.7498 |
| *T* ano. month prev. × *P* ano. month prev. | -0.0006 | 0.0003 | **0.0419** | -0.0001 | 0.0003 | 0.7196 |
| Year | not included |  |  | -0.0411 | 0.0048 | **0.0000** |
| R<sup>2</sup> | 0.6543 |  |  | 0.6661 |  |  |
| AIC | 13156.2570 |  |  | 13101.6760 |  |  |

*T*, temperature; *P*, precipitation; ano., anomalies; cur, year of sampling; prev., month of the sampling day but in the previous year; Stde, Standard Error. Bold p-values highlight significant correlations (p-value < 0.05) and the brightness of color for the “Estimate” column is proportional to the magnitude of the estimates (red for negative and blue for positive correlations).

### Contributions of weather, habitat conditions and agricultural intensification to long-term trends in insect biomass

Another important conclusion of Müller *et al*. is that weather conditions are the main drivers of the temporal changes in insect biomass, in contrast to temporal changes in habitat conditions, which play a minor role.

Since weather conditions, as well as insect biomass, exhibit strong inter-annual variations, weather conditions could drive inter-annual variability in insect biomass without being the main driver of the long-term negative linear temporal trend observed by Hallmann *et al*.^2^. In contrast, the habitat conditions chosen for the analysis (number of trees, proportion of arable land in a 200m radius, etc.) are unlikely to exhibit strong interannual variations, and thus to explain inter-annual variability in insect biomass, but could be an important driver of the long-term trend. Besides, some habitat variables extracted from Hallmann *et al*. had coarse temporal resolution. For example, proportion of habitats within the 200m radius have been calculated from two sets of aerial images, taken in 1989–1994 and 2012–2015, and yearly values have been linearly interpolated.

Available data on agricultural intensification over space and time, even if increasing, remain very scarce. To test our assumption that agricultural intensification, especially insecticide use, could be one of the drivers of the insect biomass decline observed by Hallman et *al*., we looked for quantitative proxy of agricultural intensity available over space and time. The annual yield of sugar beetroot at district level^9^, once corrected by inter-annual variation due to weather conditions, is a good candidate for a proxy of the intensity of agricultural practices, as it was until very recently systematically associated with neonicotinoid-treated seeds^10,11^ and involve fertilization, herbicides and fungicide treatments^10^. Sugar beetroot is especially prevalent in region studied by Hallman *et al*.^9^, and we assume the temporal trends in yields in a given district, corrected by local climatic conditions, relates on the agricultural intensification dynamics.

We added this spatio-temporal variable, as a proxy for agricultural intensity, to our model, which includes a fixed year effect. This variable exhibits a strong correlation with time (*r* = 0.89), but also a spatial heterogeneity that help to disentangle its link with insect biomass from the overall remaining temporal trend. To better model spatial heterogeneity in landscape sampled plots due to other anthropogenic drivers, we also included the Human Footprint index value associated with each plot, calculated over 1995-2004 at 1km^2^ scale^12^.

The result of that model shows that this proxy of agricultural intensity is the main correlates of the decline in insect biomass (Fig. 2), after the remaining temporal trend that relates to a decline due to unknown mechanisms. In parallel, the spatial variable of the Human Footprint Index (HFI), also exhibited a strong negative effects (*β* = -0.006 g.day^-1^/HFI point, p-value = 1.7e^-12^) on insect biomass, consistently with previous results^13^.

**Fig. 2:**
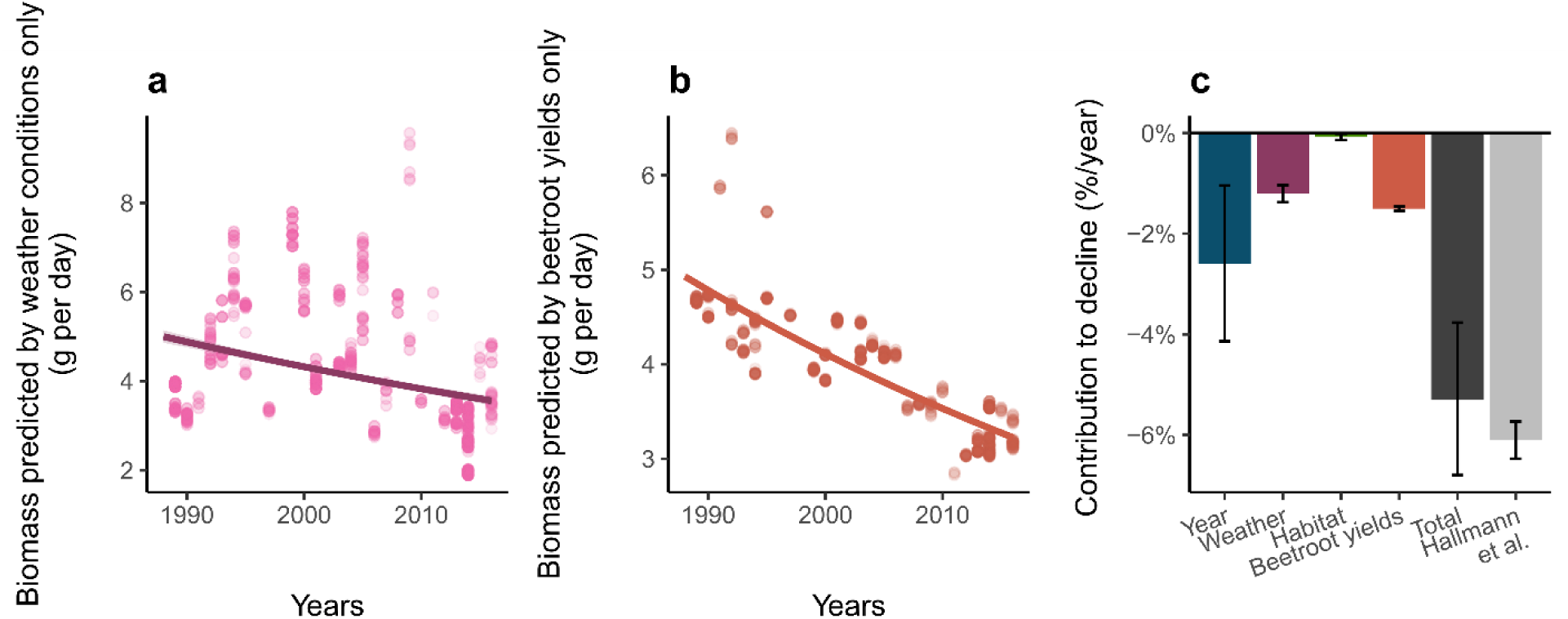
contributions of temporal changes in agricultural intensity to decline in insect biomass. (a) Biomass values predicted by (a) weather conditions and (b) agricultural intensity only, as a function of the time. The temporal trend in those values (line) is the contribution of those conditions to the long-term temporal trend in insect biomass. (c) Contributions in long-term insect decline of the remaining temporal trend (unknown mechanism), weather and habitats conditions (as modelled by Müller et al.) and a proxy of agricultural intensity (beetroot annual yields per district of the sampling location). The sum of these contribution (Total) is highly consistent with the one estimated by Hallman et al. (2017).

We are aware that this model used a too indirect proxy of agricultural intensity, which exhibits a high collinearity with time, to draw conclusions. However, it shows that when this proxy of agricultural intensity is “competing” with other drivers to explain variation in insect biomass, agricultural intensity appears to be one of the main driver of long-term insect decline, which contradicts the conclusions of Müller *et al*.^1^, but tend to support previous claim^2,6,14,15^.

## Conclusions

In writing this comment, we do not intend to tone down the effects of weather conditions on insect biomass; they are clearly demonstrated by Müller al.’s analysis, and have been supported by other studies^16–22^. The analyses of Müller *et al*. show that weather conditions could partially drive the decline in insect biomass observed by Hallmann *et al*.. However, their analyses are not suited to conclude that weather conditions are the main driver of the observed decline, or that habitats conditions contributed little to this decline. Alternative analyses even show the opposite pattern: habitats conditions play a significant role in insect biomass loss, but most of the temporal decline in insect biomass remains unexplained.

We would like here to recall that our ability to model complex ecological changes remains limited. Müller *et al*. push in the right direction in trying to understand the drivers of the temporal decline in insect biomass using correlates on the basis of extensive theory on the causal mechanisms. The lack of data about insect biomass drivers with appropriate spatial and temporal resolution is likely to be common, as it was the case for these analyses: however, not accounting for missing predictors is likely to produce highly biased results.

We also would like to warn against sweeping interpretation of row data. Müller *et al*. observed that adding recently collected data (2016-2022) to Hallmann *et al*. time series, results in a non-significant decline in biomass between 1989 and 2022 based on their Figure 1. The 1989-2016 data used by Müller *et al*. to fit their model were mostly collected in middle-west Germany, while the 2016-2022 data, used to validate the model, were collected in south-east Germany (see their Extended Data Fig. 1), an area with a smaller human footprint index than the area in which the data of Hallmann *et al*. were collected. Thus, these data do not constitute a homogenous time series and cannot be used, without statistical treatment, for interpretation about the continuation or the slowing down of the insect plight in Europe.

To understand further how insect biomass vary over time, we need models that simultaneously include all potential drivers in a similar way. Since most of the global change drivers likely depend on each other, e.g. the effect of climate change on insect abundance is mediated by land use^17^, and because data are about the spatio-temporal dynamics of these drivers are often missing, this remains a challenging task. We thus stress the need to be conservative in the interpretation of results when analysing large scale ecological patterns, to prevent over-interpretation of analyses. Drawing conclusions that are not properly supported by statistical findings is likely to disrupt both the scientific debate and public outreach, with possible negative consequences for the trust in scientific results on important topics for societies such as environmental issues.

## Acknowledgement

We thank the authors of the original study, Müller *et al*., for providing R codes and data allowing a good reproducibility of their analyses, and for their comprehensive answer. We also thank Emmanuelle Porcher, Benoit Fontaine, Ignasi Bartomeus, Grégoire Loïs and Mathilde Vimont for their helpful comments on this manuscript. FD was funded by the European Research Council (ERC) under the European Union’s Horizon 2020 research and innovation program (grant agreement N° 787638, led by C. H. Graham) and by a SNF *Postdoc Mobility* grant (P500PB_217801).

